# Investigating the Effects of Coding Variants on Protein Subcellular Localization in Autism Spectrum Disorder

**DOI:** 10.64898/2026.11.26.769325

**Authors:** Katrin Donetski, Alan Packer, Natalie Sauerwald

**Affiliations:** Department of Biomedical Engineering, Cornell University; Simons Foundation Autism Research Initiative, Simons Foundation, New York, NY, USA; Center for Computational Biology, Flatiron Institute, New York, NY, USA

## Abstract

Though many genetic variants have been associated with autism spectrum disorder (ASD), the functional effects of many of these variants remain poorly understood. Protein mislocalization, in which proteins fail to reach their intended destinations within the cell, may be one mechanism through which genetic variants contribute to cellular dysfunction. Here, we applied ProtGPS, a protein language model, to predict changes in protein localization across 12 subcellular condensate compartments for all de novo coding variants from autistic probands and their non-autistic siblings in the SPARK cohort. ProtGPS predicted small differences in localization scores for most variants, with a subset showing substantially larger changes. The fraction of variants with higher predicted localization effects varied by consequence class, with higher fractions among start-lost, frameshift, and stop-gained variants. Comparisons between probands and siblings did not reveal widespread statistically significant differences. However, significant differences were observed for missense variants affecting RNA granule localization predictions and stop-gained variants affecting nucleolus localization predictions. These results support further investigation of whether de novo variants associated with autism cause dysregulation through disruption of protein localization to biomolecular condensates. We release the predictions publicly as a resource to researchers considering these questions.

## Introduction

Autism Spectrum Disorder (ASD) is a multifaceted neurodevelopmental condition defined by the DSM-5 as having deficiencies in social communication and repetition of certain behaviors [1]. ASD is highly heterogeneous, both genetically and phenotypically, which makes it a particularly difficult condition to understand. The broad range of its physical manifestations and considerable overlap with other disorders make it challenging to pinpoint the underlying biological and genetic mechanisms. Nonetheless, exome and genome sequencing of large autism cohorts has established a substantial number of genes that are likely to be associated with an autism diagnosis when disrupted by *de novo* loss-of-function or other high impact mutations [2,3,4,5,6].

One obstacle to extracting biological information from genomic variation in autism has been the fact that many missense mutations are classified as variants of uncertain significance (VUS), both in autism and other conditions [6,7,8]. Although significant efforts are underway to classify autism VUS as either pathogenic or benign [9,10,11,12], many more variants remain unassessed, leaving a significant number of individuals without a definitive genetic diagnosis. Moreover, the range of functional effects that have been tested remains somewhat restricted. For example there is a growing body of literature on variants that disrupt the functions of biomolecular condensates [13], which are membrane-free protein assemblies with a unique chemical environment formed through phase separation. This organization is driven by the weak and flexible interactions of intrinsically disordered regions in each protein.

Disruptions of condensate formation are of particular interest in the context of autism in light of computational predictions that autism-associated proteins tend to be enriched for the biophysical properties that are involved in phase separation [14]. In addition to computational predictions made en masse, there are compelling individual examples of autism-associated mutations disrupting condensate function. For example, abnormal splicing in the neuronal gene CPEB4 has been linked to autism, and CPEB4 forms biomolecular condensates [15]. CPEB4 is an RNA-binding protein that acts by elongating poly(A) tails to promote mRNA translation. There is a neuron-specific microexon within CPEB4 that has been shown to interact with histidine residues, consequently preventing the histidines from permanently clustering with each other. When the alternative splicing pattern is disrupted, there is decreased inclusion of the microexons in the CPEB4 proteins. The microexons are crucial for binding to the histidine clusters to form reversible condensates, otherwise the histidines irreversibly bind to each other, forming a permanent bond. The reversible nature of these condensates is important for dissolution once the neurons are stimulated. Upon neuron depolarization, a shift in the intracellular pH destabilizes interactions that form the condensates, allowing for CPEB4 to release and activate translation of target genes.

In light of these and other data, there is a need for additional computational and functional assessments of autism-associated VUS and their possible roles in biological condensate formation. ProtGPS is a protein language model that predicts where proteins are likely to localize within the different condensates in the cell [16]. It is an adaptation of the ESM2 protein language model with a neural network classifier, and offers a new approach to the prediction of protein behavior. The model takes a linear protein sequence as an input, and produces 12 independent probabilities of this protein being found within each of the 12 different condensate compartments. The model is trained with the following condensate compartments: RNA granule, cell junction, stress granule, nucleolus, nuclear pore complex (NPC), cajal body, PML body, transcriptional condensate, nuclear speckles, chromatin, post synaptic density (PSD, and P-body. Here we apply ProtGPS to a body of autism genetic data to gain new insight into the fraction of autism-associated VUS that might be classified as pathogenic owing to their roles in modulating biological condensate function. We provide the predictions as a resource to other researchers exploring potential variant effects in autism.

## Methods

### Data

We analyzed whole-exome sequencing data from SPARK, including autistic probands and their non-autistic siblings [17,18]. The cohort described in this draft comprises 13,324 probands and 5,334 siblings with complete child-parent trio data. De novo coding variants (DNVs) were defined as variants present in the child and absent from both parents [19,20].

DNVs were identified using the HAT pipeline [21], with DeepVariant (v.1.1.0) [22] and GATK HaplotypeCaller (v.4.1.2.0) [23]. Candidate variants were required to be identified by both callers as present in the child and absent from either parent. Subsequent filtering excluded low-confidence calls, including variants in recent repeats, low-complexity regions, and other unreliable genomic regions.

Variants were annotated using Ensembl Variant Effect Predictor (VEP release 111.0), with LOFTEE (v.1.0.4) and AlphaMissense plugins [24–26]. We retained variants annotated as missense, in-frame insertion or deletion, protein-altering, stop-gained, frameshift, splice-donor or splice-acceptor, start-lost, stop-lost, or transcript ablation. Intronic and untranslated-region variants were excluded from the protein-sequence analysis.

### Reference and variant protein sequences

For each variant-transcript pair, we obtained a reference transcript sequence and generated a variant transcript by introducing the variant at the corresponding position. Variants were analyzed independently, so each altered transcript contained a single introduced variant. To account for alternative splicing, we considered all known gene isoforms represented in the GENCODE v47 basic transcript annotation for GRCh38 (hg38). Reference and variant transcripts were translated into amino acid sequences using the Biotite Python package [27].

### ProtGPS predictions and localization effect metrics

We applied ProtGPS [16] separately to each reference and variant protein sequence to obtain 12 independent localization probabilities for each sequence. For each compartment, the signed prediction difference was defined as the variant probability minus the reference probability; its absolute value summarized the magnitude of the predicted change. A difference less than zero indicated reduced predicted localization to that compartment, and a positive difference indicated increased predicted localization. Because the 12 outputs are independent, a decrease in one compartment does not by itself imply redistribution to another.

The maximum localization effect was defined as the largest absolute prediction difference across the 12 compartments for a reference–variant protein pair. We used a maximum difference greater than 0.5 to identify variants with higher predicted localization effects for descriptive characterization (Figure 2). This threshold defines a prediction-based subset and does not establish pathogenicity or experimental disruption of condensates.

### Variant characterization and proband-sibling comparisons

We summarized variant consequences in the full analysis set (Figure 1A) and the distributions of absolute localization prediction differences by consequence (Figure 1B). For each consequence, the fraction with higher predicted effects was calculated as the number exceeding the maximum-effect threshold divided by the total number in that consequence (Figure 2A). These percentages have separate denominators and are not expected to sum to 100%. We also characterized the compartments affected within the higher-effect subset (Figure 2B).

**Figure 1.**
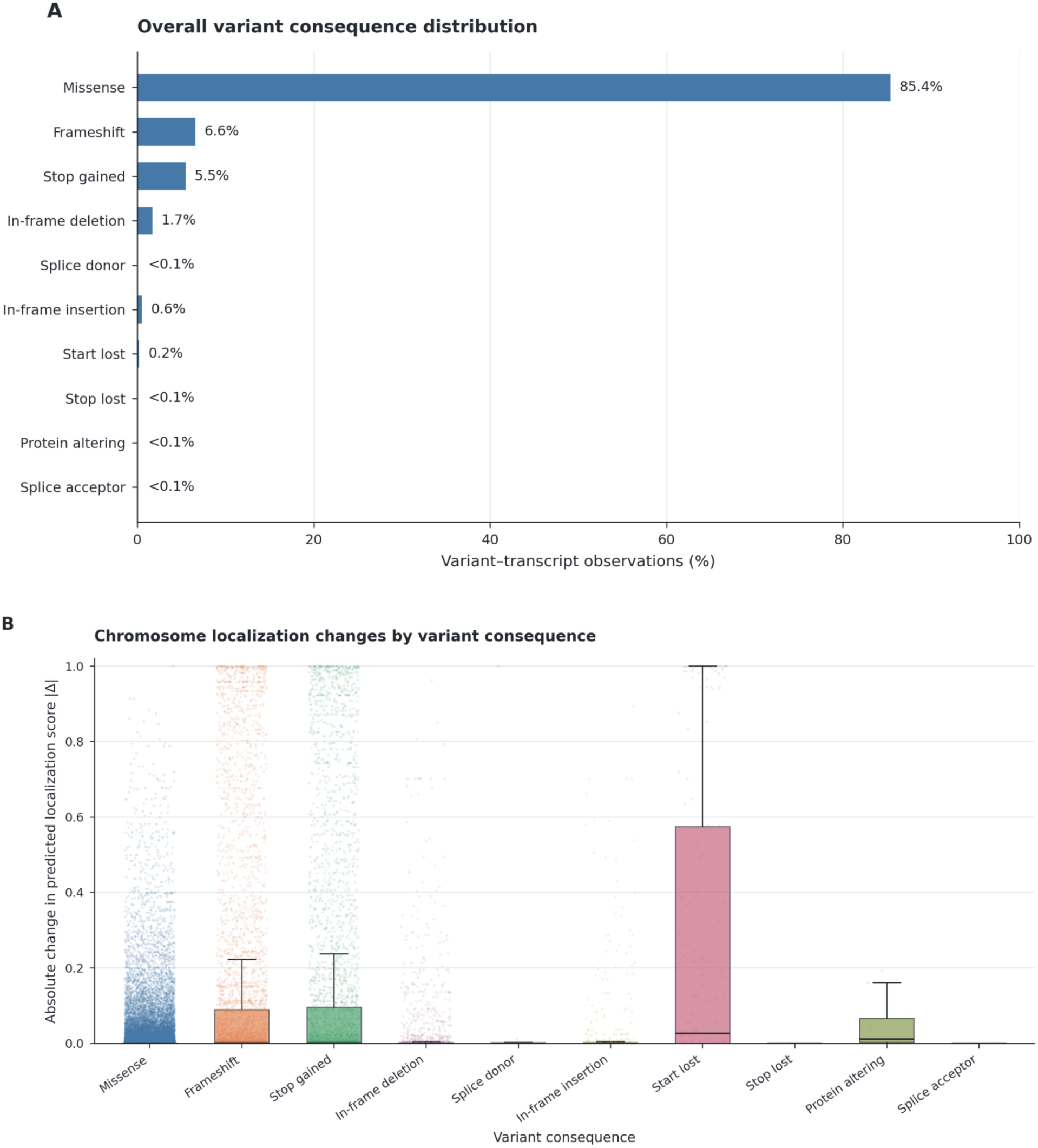
**A.Variant consequence distribution and predicted localization changes.** Overall distribution of the variant consequences, shown as a percentage of variants across all analyzed transcripts. Missense variants are the majority, with frameshift and stop-gained variants following. **B. Absolute changes in predicted chromosome condensate localization across variant consequences**. Each point shows the absolute difference in the predicted chromosome localization score between the reference and variant protein sequences for the same transcript. These differences are grouped by variant consequence, such that each box summarizes the distribution of the absolute predicted differences. The line inside each box marks the median, and the box edges mark the lower and upper quartiles. The whiskers extend to the smallest and largest observed differences that are within 1.5 times the interquartile range below the lower quartile and above the upper quartile, respectively.

**Figure 2.**
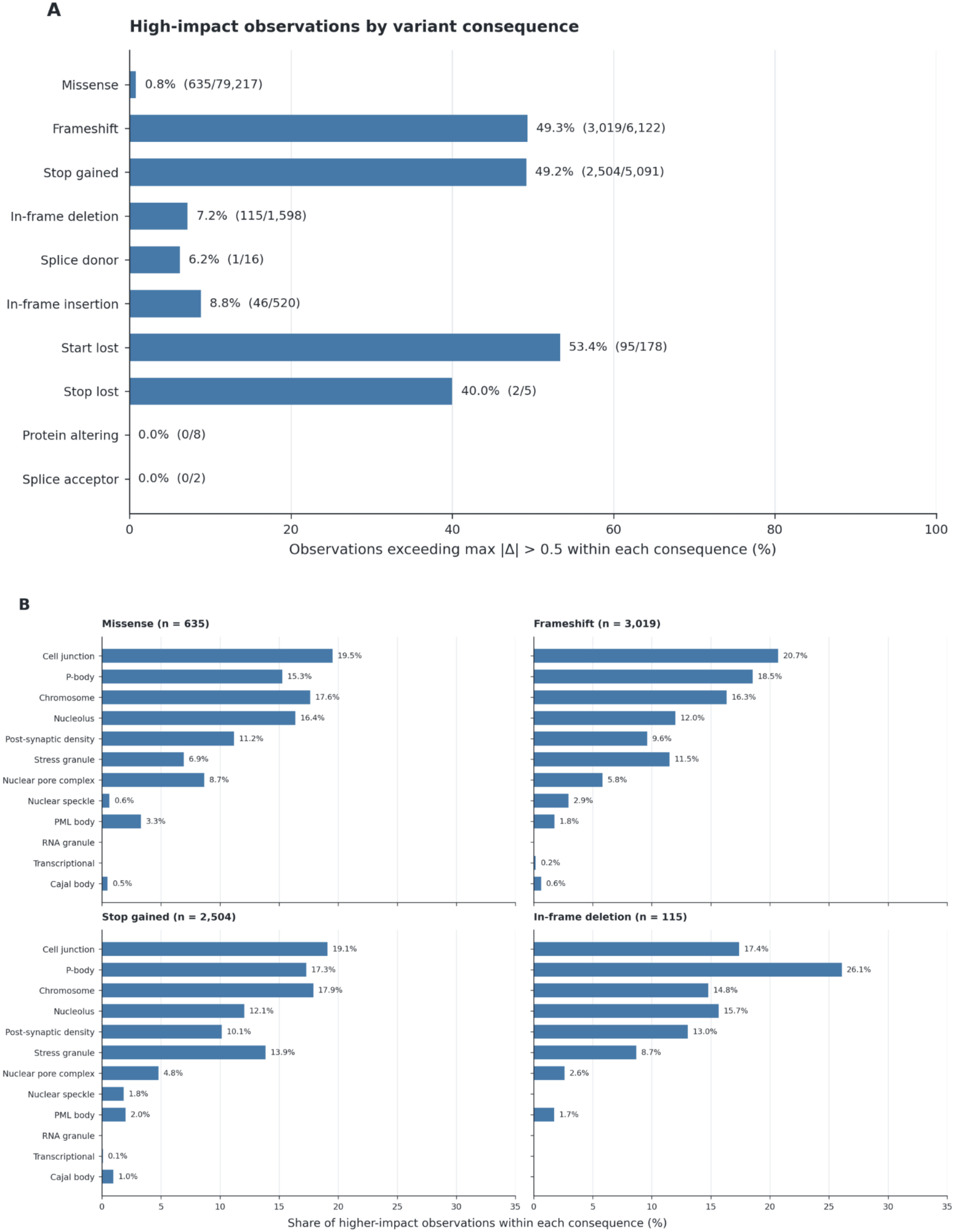
Characterization of variants with higher predicted localization effects. **A**. Percentage of variants across analyzed transcripts with a maximum absolute predicted localization difference greater than 0.5 within each consequence class. Percentage labels show the number exceeding the threshold divided by the total analyzed within that class. **B**. Distribution of compartments with the largest absolute predicted localization difference among variants exceeding the threshold. Percentages are calculated separately within each of the four consequence classes shown.

To compare probands and siblings, we examined group-stratified distributions of absolute prediction differences (Figure 3A). For each compartment–consequence combination, Welch’s t-test was used to compare mean differences in predicted localization scores (variant score − reference score) between groups. Benjamini–Hochberg correction was applied across all valid tests to control the false discovery rate (FDR), and results were summarized as −log10(q) in a heatmap (Figure 3B). These tests compare prediction differences, rather than frequencies of variants.

**Figure 3.**
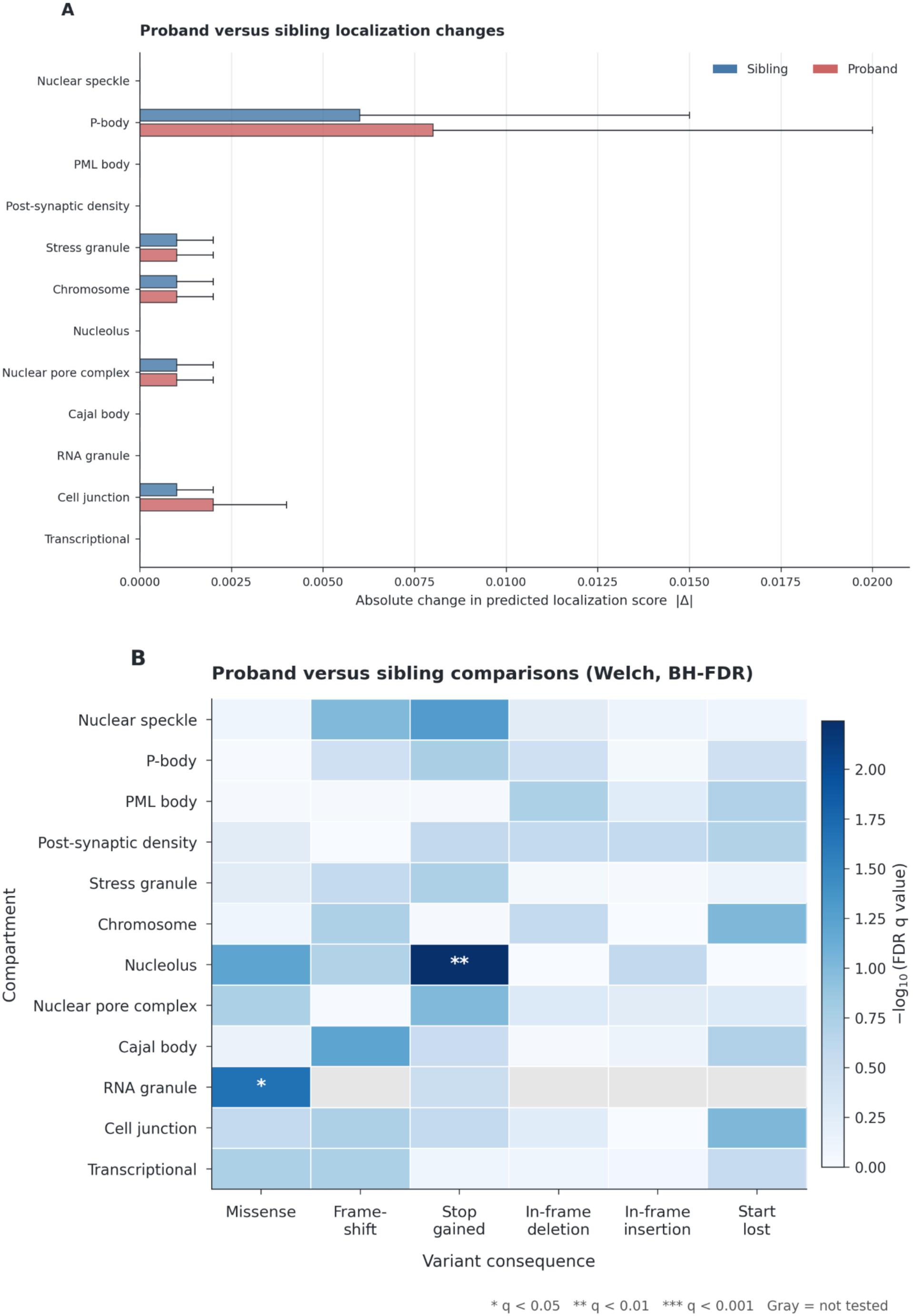
Comparison of localization effects between probands and siblings across compartments. **A**. The boxplots show the distributions of absolute localization prediction differences for all 12 compartments, with variant consequence types combined. **B**. Statistical comparison of predicted localization effects between probands and siblings. The heatmap shows the significance values from Welch’s t-test comparing mean localization prediction differences (variant score–reference score) between groups after Benjamini–Hochberg correction. The t-test was performed individually on each of the condensate compartment and variant consequence class combinations. The color on the map represents the -log10 (q-value), with stars indicating significance (* q < 0.05, ** q < 0.01, and *** q < 0.001). Gray cells indicate comparisons that were not tested. Significant differences were observed for the missense variants affecting the localization predictions in the RNA granules and stop-gained variants affecting nucleolus predictions.

## Results

### Variant landscape and overall predicted localization effects

The SPARK dataset we analyzed here contained 29,029 unique coding DNVs mapping to 36,789 transcripts across 11,508 genes, with an average of 1.56 coding DNVs per proband and 1.52 per sibling. Missense variants accounted for 85.4% of coding DNVs, followed by frameshift variants (6.6%) and stop-gained variants (5.5%) (Figure 1A).

ProtGPS predicted small absolute changes in localization probability scores for most observations, with a subset showing substantially larger changes (Supplementary Figure S1). Plotting the absolute difference in predicted localization probability between reference and variant sequences suggests that the majority of variants likely have little effect on protein localization, with score differences at or near zero. However, there are a number of variants with very high predicted localization effects within the chromosome condensate (Figure 1B). In particular, while the median absolute differences for the largest categories, missense, frameshift, and stop-gained variants, are extremely small, all three categories show a high number of variants with localization differences greater than 0.5, suggesting many of these could have a significant effect. Start-lost variants had the widest interquartile range of the predicted chromosome localization changes (Fig. 1B). Large changes for frameshift and stop-gained variants were also seen in several other compartments, including P-body and cell junction (Supplementary Fig. S1). An early stop codon can truncate the protein sequence, and a frameshift can substantially alter the sequence downstream of the variant. These changes may help explain some of the large differences in predicted localization scores.

### Characterization of variants with higher predicted localization effects

We next focused on variants with a maximum absolute localization prediction difference greater than 0.5 to characterize the consequences and compartments associated with larger predicted effects. Of the 92,757 analyzed variants across transcripts, 6,417 (6.9%) exceeded this threshold and were classified as being higher-impact variants. The fraction of high-impact variants varied by consequence class, including 53.4% of start-lost variants, 49.3% of frameshift variants, 49.2% of stop-gained variants, and 0.8% of missense variants (Figure 2A).

Among the high-impact variants, cell junction was most frequently associated with the largest predicted localization changes for missense (19.5%), frameshift (20.7%), and stop-gained variants (19.1%) . For in-frame deletions, P-body was most frequently associated with the largest predicted localization change (26.1%; Figure 2B).

### Comparison of predicted localization effects between probands and siblings

We compared the absolute differences in predicted localization scores between reference and variant protein sequences in probands and siblings across all 12 compartments. Most compartments had similar boxplot summaries for both probands and siblings, with median differences close to zero. P-body showed the widest interquartile range in both groups, with a wider range in probands than siblings. Cell junction also showed a wider interquartile range in probands (Figure 3A).

The compartment-specific tests did not identify widespread statistically significant differences between probands and siblings. Two of the compartment-consequence combinations were reported as significant after correction: missense variants for RNA granule predictions and stop-gained variants for nucleolus predictions (Figure 3B). Thus, the reported differences were limited to specific combinations rather than a broad shift across compartments and consequence classes.

## Conclusion

Overall, these results support further analysis of whether de novo variants associated with autism disrupt protein localization to biomolecular condensates. While this investigation did not reveal broad differences between probands and siblings, there were specific combinations of condensates and variants that revealed a significant difference. These preliminary differences suggest that condensate disruption affecting specific mechanisms, particularly within certain compartments and variant classes, may contribute to autism risk and requires further investigation. Future work could validate whether these patterns are reproduced by other protein localization prediction tools and evaluate effects produced by combinations of variants, which are missed when each variant is analyzed independently. Additionally, future investigation is needed to explore whether variants within intrinsically disordered regions are more likely to alter predicted protein localization. The functional effects of many variants of uncertain significance remain poorly understood and difficult to characterize. These predictions may also help prioritize such variants for functional assessment in autism.

RNA-binding proteins, including the aforementioned CPEB4, are critical in maintaining cellular homeostasis. The impact of dysregulation at the level of both the RNA-binding proteins and their downstream target sites affect neuropathogenic pathways in ways that could drive psychiatric disorders, including autism [28,29]. Variants associated with an RNA helicase, encoded by the DDX3X gene, are a leading cause of ASD [30]. The helicase functions by regulating RNA metabolism, involving RNA-containing condensates and granules that form via phase separation. As yet another example, the RNA-binding protein SFPQ forms meshwork-like biological condensates and is involved in the transcriptional elongation and splicing of extra-long genes, many of which are associated with autism [31]. Thus, the data implicate abnormal RNA granule assembly, potentially driven by mislocalization, as a mechanism of autism pathogenesis.

In the specific chromosomal compartment, there are higher predicted effects in probands than their healthy siblings. This may be relevant to the large number of chromatin-associated proteins known to be associated with autism likelihood..

Targeting condensate dysfunction offers a potential therapeutic approach, as shown by zinc supplementation restoring proper condensate formation to mitigate the effects from ASD-linked mutations [32]. Other approaches could involve correcting splicing patterns or directly manipulating genetic variants to counteract disruptions to condensate regulation. Being that certain genes are more prone to condensate dysfunction, identifying these risk factors could reveal underlying mechanisms of disease that act on a molecular level. Genetic variants can drive mislocalization of proteins following disease-specific patterns, emphasizing the potential to identify and correct such disruption at the cellular level.

## Data Availability

To abide by the informed consent that individuals with autism and their family members signed when agreeing to participate in a SFARI cohort (SSC and SPARK), researchers must be approved by SFARI Base (https://base.sfari.org).

## Code Availability

Code and scripts used for the analyses presented in this manuscript are available via GitHub at https://github.com/Kdonetski/ASD-protgps.

