## Supplemental Figure S1 for "Investigating the Effects of Coding Variants on Protein Subcellular Localization in Autism Spectrum Disorder"

### Supplementary Figure S1

**Nuclear speckle localization changes by variant consequence**

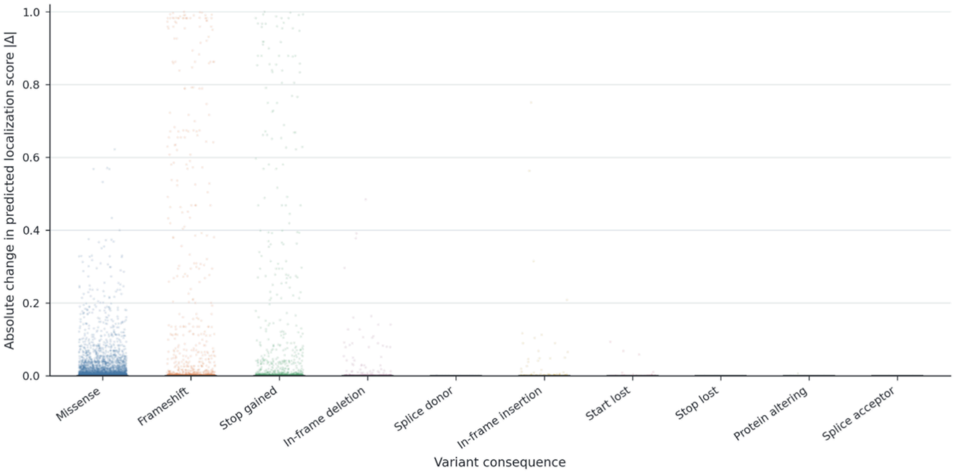


**P-body localization changes by variant consequence**

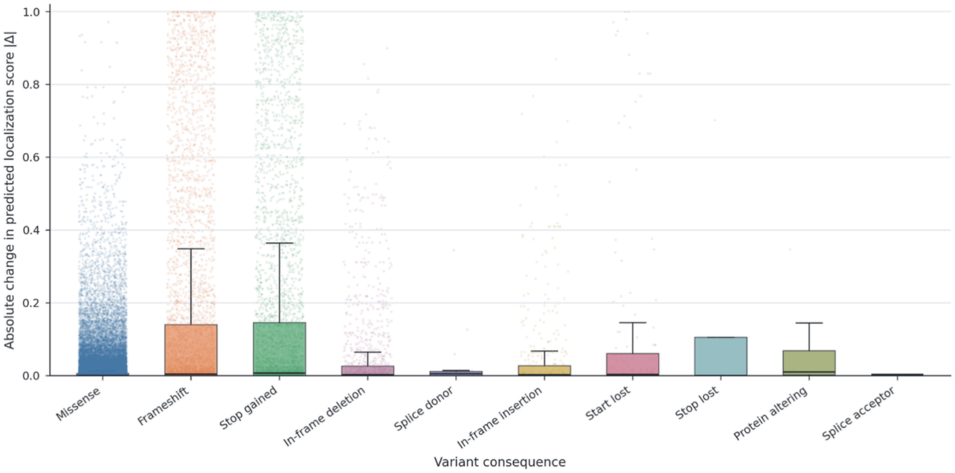


**PML body localization changes by variant consequence**

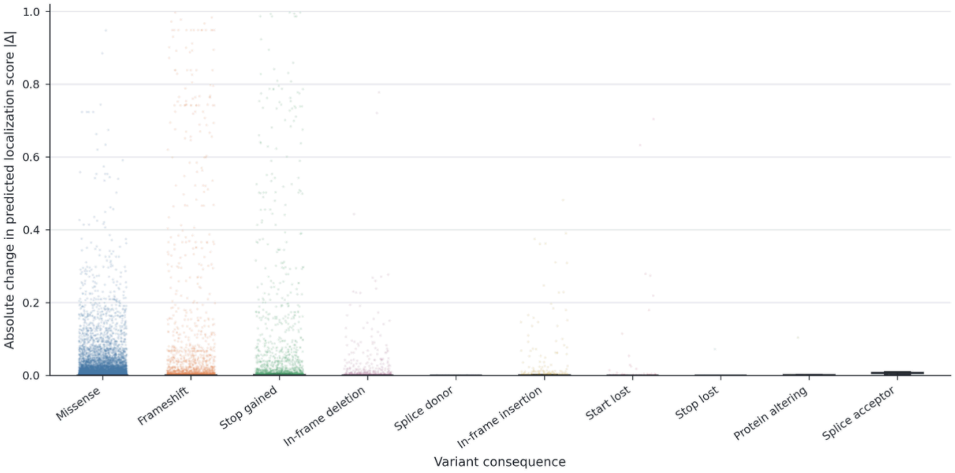

### Supplementary Figure S1

**Post-synaptic density localization changes by variant consequence**

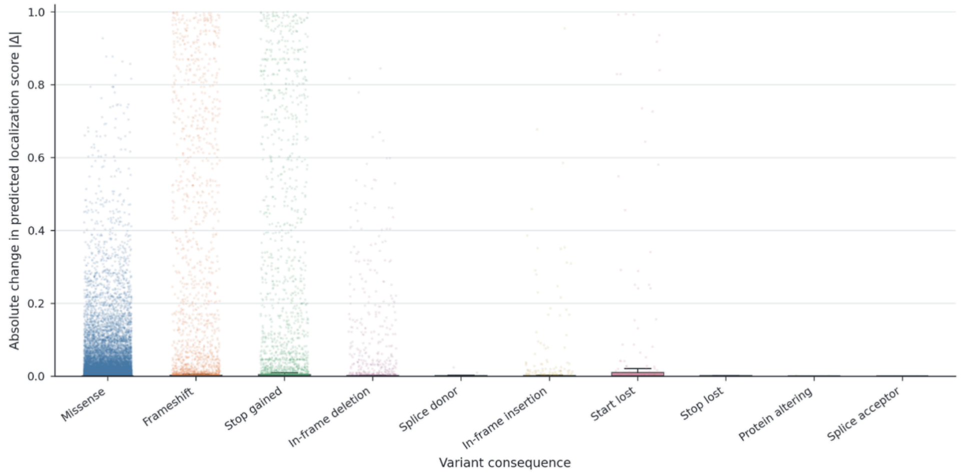


**Stress granule localization changes by variant consequence**

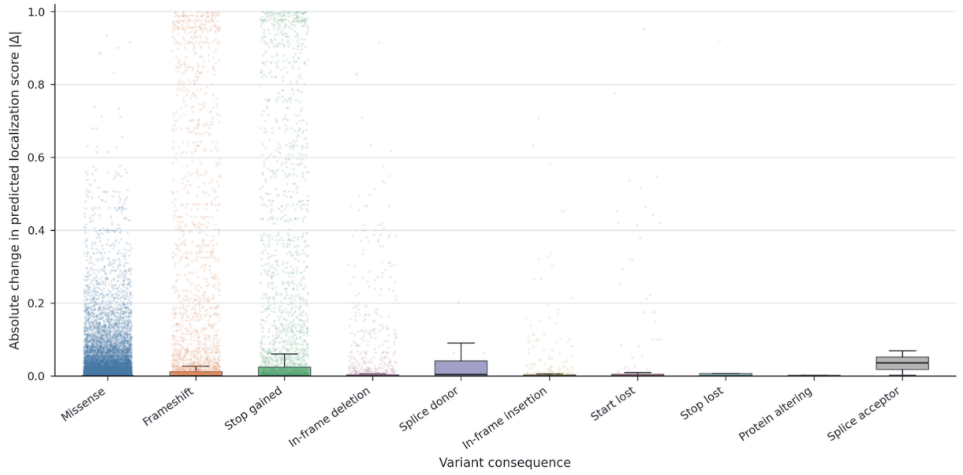


**Chromosome localization changes by variant consequence**

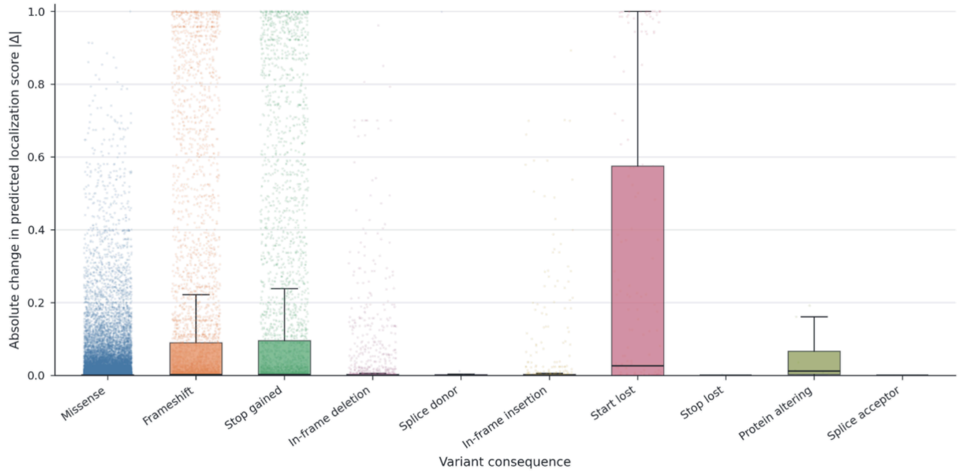

### Supplementary Figure S1

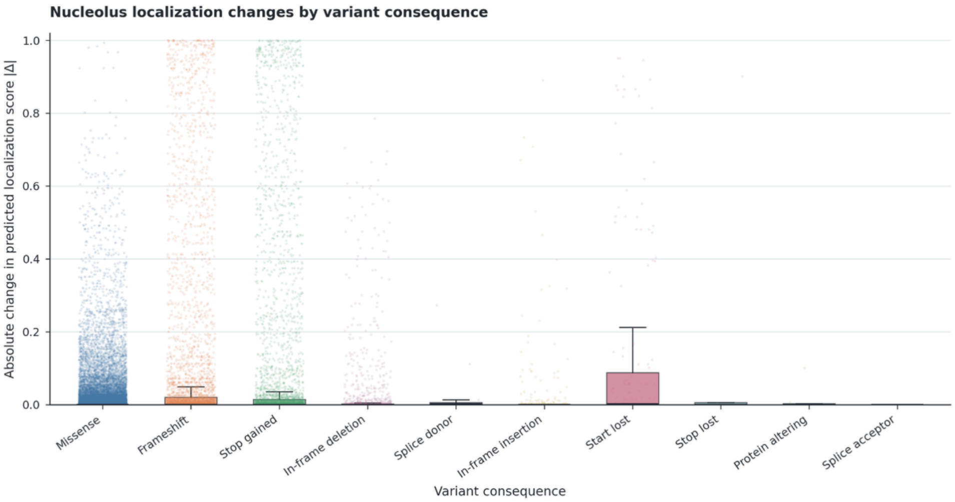


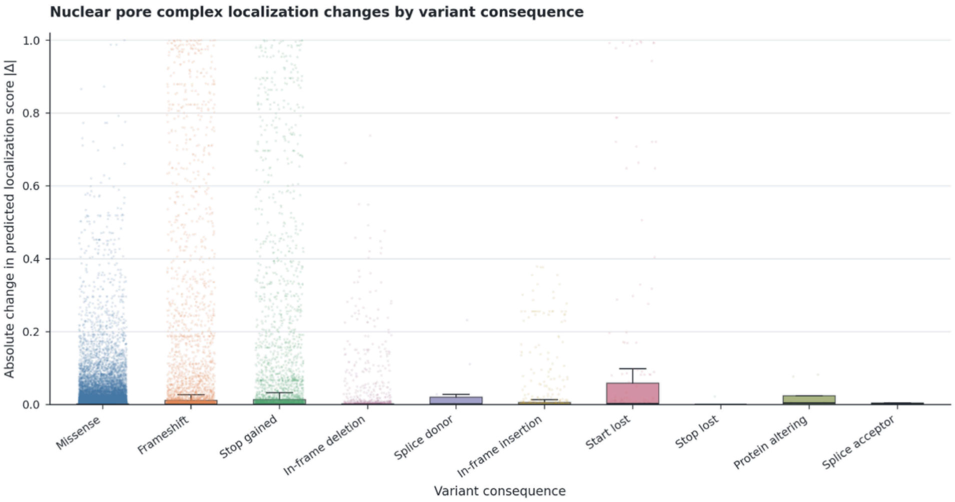


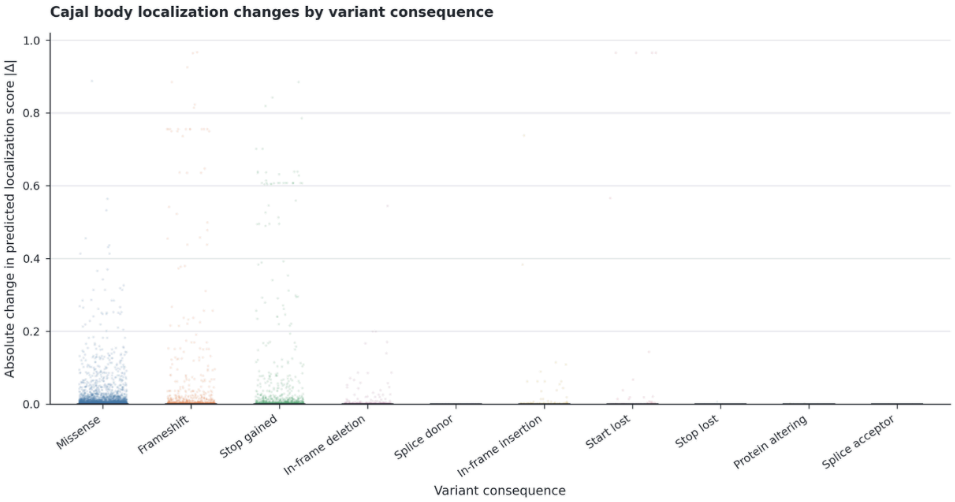

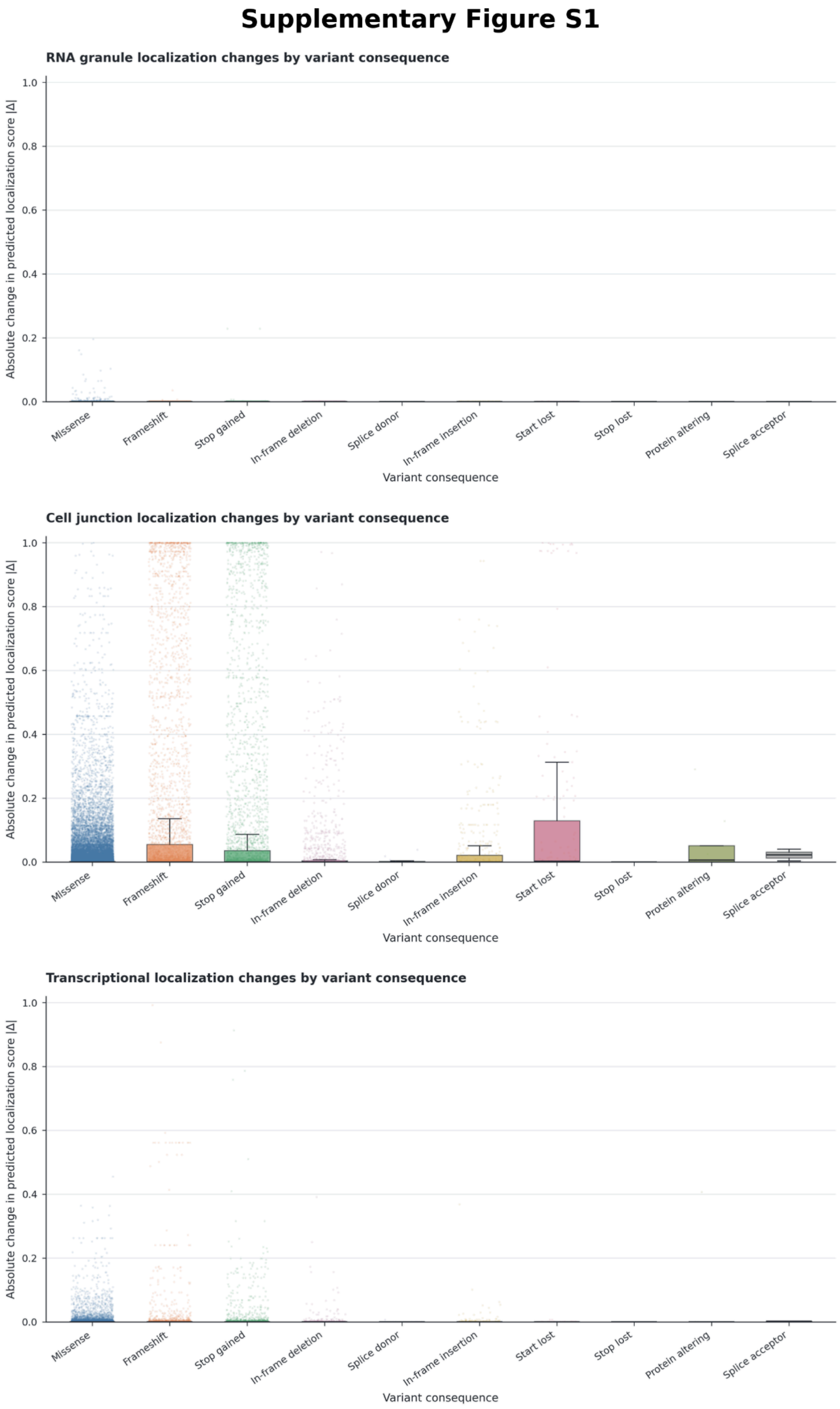


**Supplementary Figure S1. Absolute differences in predicted localization scores across all 12 compartments.** Differences between reference and variant protein scores are grouped by variant consequence. Points, boxes, median lines, and whiskers follow the conventions in Figure 1B.
